# A peculiar evolutionary feature of monkeypox virus

**DOI:** 10.1101/2022.06.18.496696

**Authors:** Yicong Dai, Xucong Teng, Difei Hu, Qiushuang Zhang, Jinghong Li

**Affiliations:** Department of Chemistry, Center for BioAnalytical Chemistry, Key Laboratory of Bioorganic Phosphorus Chemistry & Chemical Biology, Tsinghua University, Beijing, China

**Keywords:** monkeypox virus, translation regulation, genomic evolution, RNA secondary structure

## Abstract

From 1 Jan 2022 to 15 June 2022, more than 2,100 cases of monkeypox have been reported in 42 countries. This unusual outbreak of monkeypox has raised new concerns in academia and the public. To keep abreast of the trend of the monkeypox epidemic, it is extremely urgent and important to surveille the accumulated genomic mutations and the change of the transmission ability of the pathogen, monkeypox virus (MPXV). We report a non-canonical RNA secondary structure, G-quadruplex (RG4), that surprisingly evolved stepwise with various variants. The RG4 motif is located in the coding sequence region of MPXV C9L gene that is functional in inhibiting host innate immune response. The evolution decreases the stability of this RG4 and promotes C9L protein level in living cells. Importantly, all the reported MPXV genomes in 2022 contain the most unstable RG4 variant, which may be the reason of the increasing spread of MPXV. These findings recommend that health authorities and researchers pay attention to the genomic evolution of MPXV.

## Introduction

Monkeypox is caused by Monkeypox virus (MPXV), a member of the family *Poxviridae* that is close relative to variola virus(*1*). MPXV is considered to be not very contagious and is mainly endemic in sub-Saharan Africa(*2*). Strikingly, from 1 Jan 2022 to 15 June 2022, more than 2,100 cases of monkeypox have been reported in 42 countries (**Figure 1A-1B**). Although some cases have been reported outside Africa since 2003(*3*), it is very unusual that such a large number of cases have occurred around the world. Because of the high fatality rate of 1% to 11%(*4*), the spread of MPXV may increasingly exacerbate the global health crisis. In addition to the ongoing pandemic of SARS-CoV-2, the sudden outbreak of monkeypox has raised new concerns around the world. The reason for the unusual increase in human-to-human transmission of MPXV remains a mystery(*5–7*). Therefore, it is very important and urgent to monitor the genetic features of MPXV and explore the biological mechanisms underlying the accelerated transmission of MPXV.

**Figure 1.**
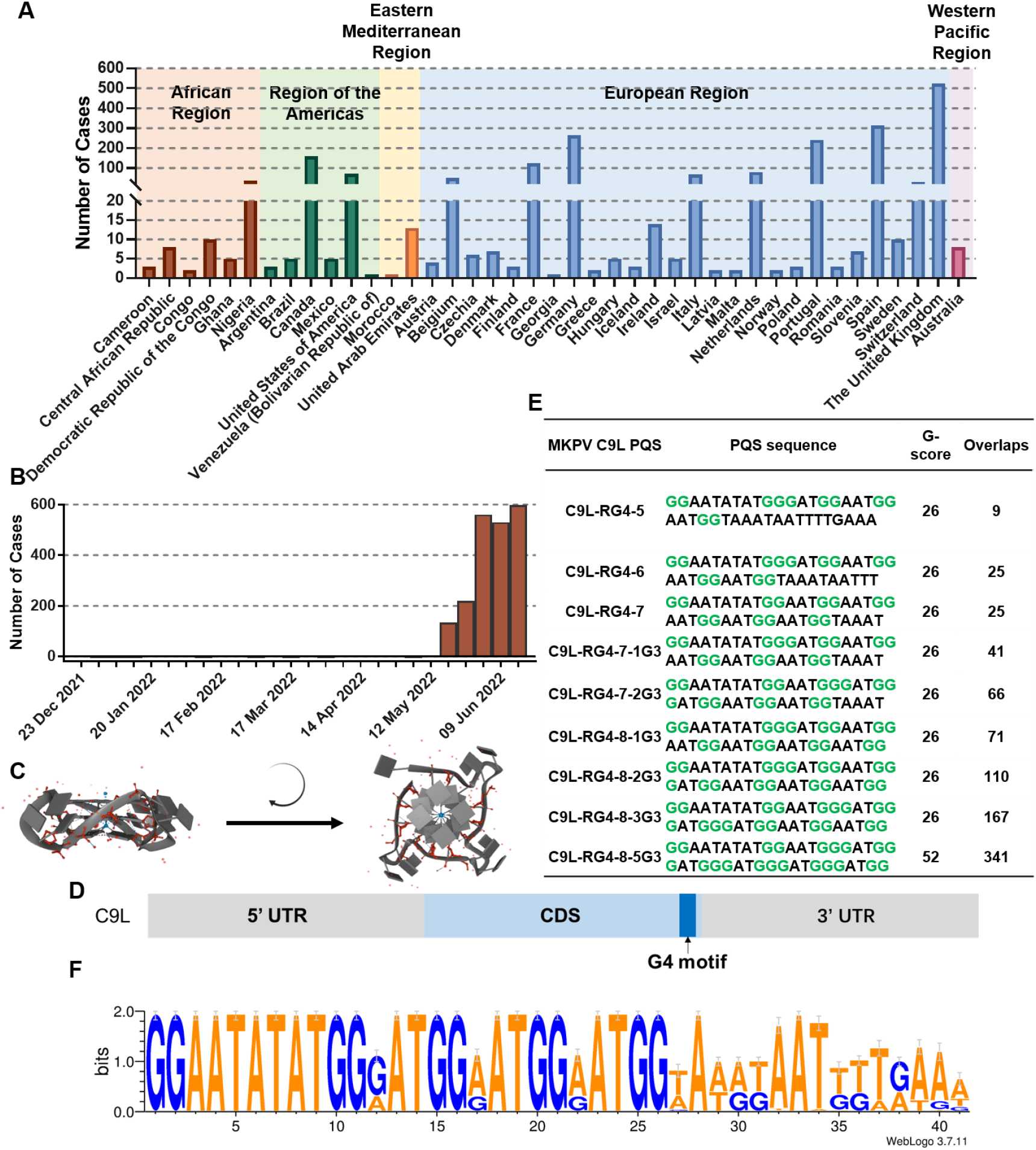
Spreading of MPXV and its RG4 motif features. **A**) Number of cases in 42 countries. Source: World Health Organization. **B**) The recent burst of MPXV case numbers across the world. Source: World Health Organization. **C**) Schematic illustration of a RG4 structure. **D**) The PQS location in the MPXV C9L gene. **E**) The sequences and features of the 9 C9L PQS variants. G-scores and overlaps are predicted by QGRS-mapper under the parameters of “Max Length: 35, Min G-Group Size: 2, Loop size: from 0 to 36”. **F**) Conservation analysis of the C9L PQS. A total of 177 complete genomic sequences of MPKV genome from the GenBank database were aligned using WebLogo 3 software.

Despite various mutation analysis of the evolving MPXV genomes, the evolution on the nucleic acid secondary structure and its impact on the viral biology are still unclear. For example, RNA G-quadruplex (RG4) is a non-canonical secondary structure of RNA(*8*). In RG4, Hoogsteen hydrogen-bonded G-tetrads stack 2-3 layers to form the quadruplex topology (**Figure 1C**). RG4 exists in many organisms, and regulates various RNA-related biological processes, including translation, translocation, RNA splicing and so *on*(*9–11*). In many viruses, such as human immunodeficiency virus (HIV)(*12*), human papillomavirus (HPV)(*18*), herpes simplex virus (HSV)(*14*), hepatitis C virus (HCV)(*15*), Epstein-Barr virus (EBV)(*16*) and SARS-COV-2(*17*), the RG4 can repress the expression of viral proteins and may have an important role in the virus life cycle(*13*). Therefore, RG4 has been proposed to become an antiviral target for therapeutic application(*19, 20*). Given that MPXV has a ~197 kb double-stranded DNA genome with 190 coding genes, it is likely that functional RG4 structures are present in the mRNA of these genes. Whether the RG4 is involved in the regulation of the biological processes of MPXV, and the relationship between the RG4 motifs and the evolution of MPXV genome, are of great interest and importance.

Herein, we first discover that there are a RG4 motif in the MPXV C9L mRNA, and it has evolved stepwise. We characterize that as the C9L RG4 motif evolves, its structural stability is gradually reduced. The MPXV strains found in the recent 5 years all have the most unstable RG4 variant. Next, we first confirm at the single-molecule level that the C9L RG4 can be formed in mammal cells. Finally, we verify that the decreased C9L RG4 stability increases the expression level of its corresponding protein. This may be the reason for the rapider transmission of the current epidemic MPXV strains, because the C9L gene of MPXV has a key role in the evasion of the innate immune response. In this way, as the MPXV further evolves, its ability to spread may continue to increase. These findings propose a new direction for investigating MPXV transmission and pathogenesis.

## Results and Discussion

We used QGRS-mapper(*21*) to analyze the putative G4-forming sequences (PQSs) in the whole MPXV genome. Given that MPKV has gradually accumulated mutations, we collected 177 complete genomic sequences of MPKV genome from the GenBank database to predict the PQSs in all the viral strains and assess the levels of sequence conservation of the PQSs. Among all the MPKV genomic sequences, there is only one PQS located in the coding sequences (CDS) region of the C9L gene (**Figure 1D**). The C9L gene of MPXV encodes a Kelch-like protein that plays an important role in inhibiting innate immune responses of the host(*22, 23*). We found that there are 5-8 G-tracts in the 9 variants of C9L PQSs (**Figure 1E**). Some C9L PQS variants has G-tracts with 3 G-bases (Blue), while others have only G-tracts with 2 G-bases (Green). Additionally, the C9L PQSs of different viral strains have diverse G-scores and overlaps, which means different G4-forming potential and stability. 5 G-tracts of the C9L PQSs are high conserved in MPXV genome (**Figure 1F**), indicating a potential function in MPXV life cycle.

To identify the evolutionary relationships between the C9L PQSs from different MPXV strains, we analyzed the phylogenetic tree based on 177 whole-genome DNA sequences of MPXV. As shown in **Figure 2 and Figure S1**, MPXV has two major genetic clades—the Central African clade and the West African clade (*27*), and some sub-clades. All the reported genomic sequences from 2022 MPXV cases are highly related to each other, and robustly belong to B.1 clade of West African clade. We then focused our analysis on the C9L RG4 feature of the strains on the phylogenetic tree. Recently reported MPXV genomes of A.1, A1.1, B.1 clade all contain C9L-RG4-5, the PQS with the minimal G-tracts, while the other clades have PQSs with more G-tracts. This finding suggests a potential evolutionary function of the C9L RG4 motif.

**Figure 2.**
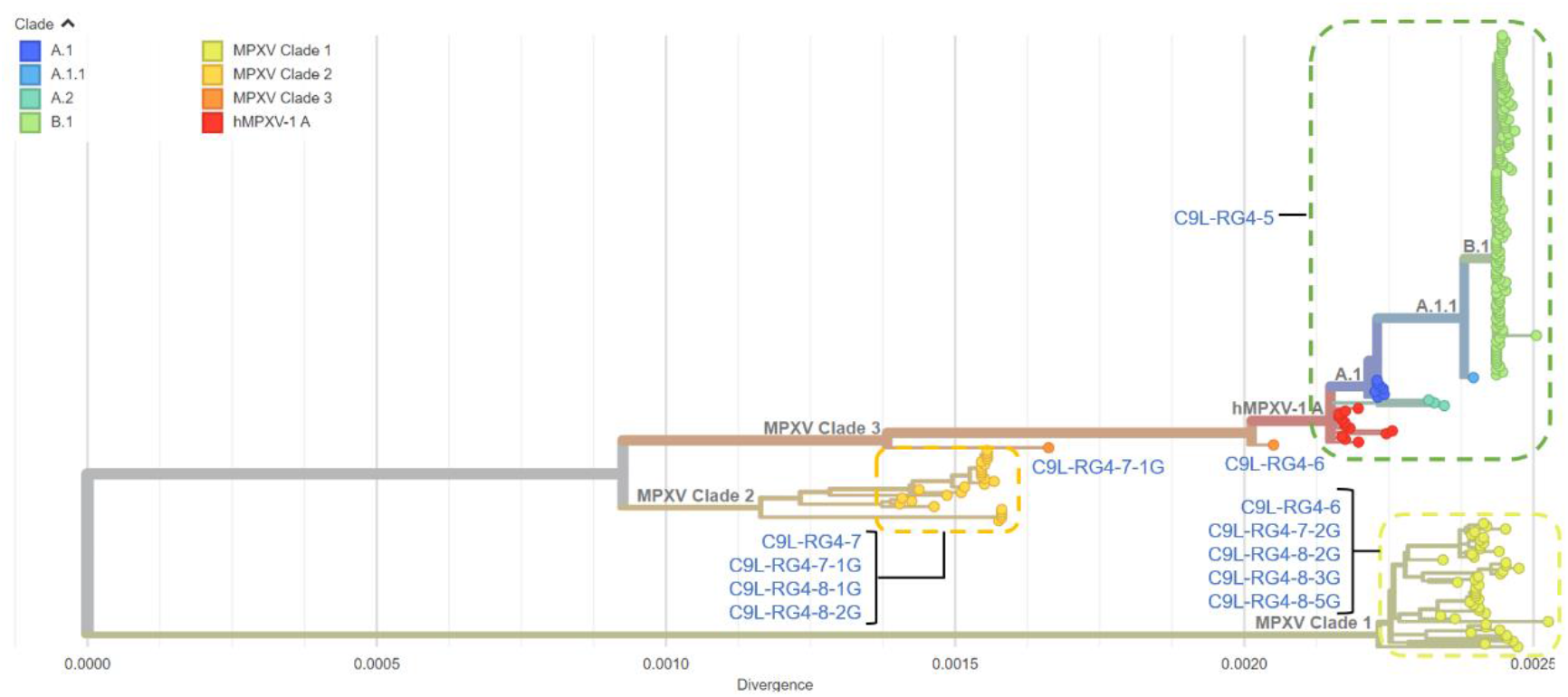
Maximum likelihood phylogenetic tree of nucleotide sequences of the wholegenome of 177 MPXV strains. The MPXV genomes were divided into 8 clades. The corresponding C9L PQS type of each clade was labelled near the clade. Downloaded and adapted from Nextstrain website.

To evaluate the formation and stability of the RG4 structures in the 9 C9L PQSs, we conducted a fluorescent turn-on assay by a widely used G4-specific fluorescent probe, thioflavin T (ThT)(*24*). After the C9L PQS oligos annealed in K^+^ containing cytosol mimic buffer, the fluorescent signal of ThT was obviously increased (**Figure S2**). In contrast, when the G-tracts of C9L PQS were G-A mutated, no fluorescence intensity increase could be observed. These results indicated that the 9 C9L PQSs can form G4 structures in a general cytosol mimic condition. Notably, at the same concentration of the C9L PQS oligos and ThT, the C9L PQS oligos generated various fluorescent intensity at the peak wavelength (**Figure 3A**), indicating that the accumulated mutations of the C9L G4 motif may regulate the stability of the RG4 structures. To further confirm the formation of G4 structures, circular dichroism (CD) measurements were carried out. As shown in **Figure S3**, the 9 C9L PQS oligos generated a positive peak near 264 nm and a negative peak near 240 nm, which is consistent with the typical CD spectra of parallel RG4 topology(*25*). When the G-tracts were G-A mutated or the C9L PQS oligos were annealed without K^+^, the corresponding CD signature disappeared, further confirming the formation of G4 structures. In addition, the melting temperature (*Tm*) of the 9 C9L PQS oligos was measured by detecting the temperature-dependent change of the CD signals at 264 nm. The *Tm* values of C9L PQS oligos varied from 60.3 to 85.5 °C (**Figure 3B**). As we anticipated, the thermal stability of G4 structures formed by different C9L PQSs was significantly different, which was in agreement with the bioinformatical prediction.

**Figure 3.**
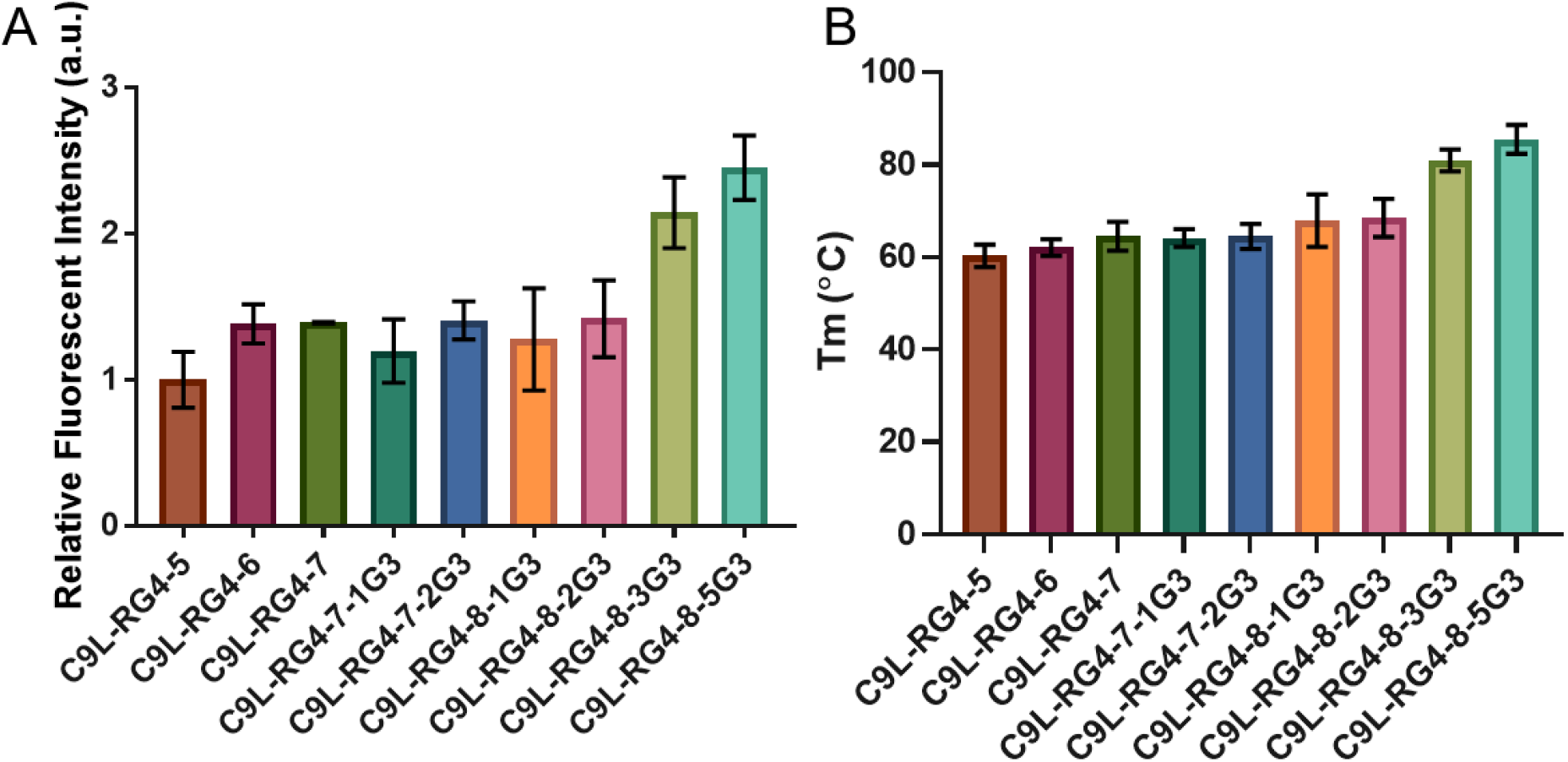
Stability of the C9L RG4 motifs. **A**) Fluorescence intensity of the 9 C9L PQS oligos at the peak wavelength in ThT fluorescence turn-on assay. **B**) Melting temperatures of the 9 C9L PQS oligos measured by temperature-dependent CD spectra. Data are shown as mean ± SEM of three independent experiments.

We next sought to determine whether the 9 C9L PQSs could fold into RG4 structures in living cells. Since live virus culture requires very strict biosafety laboratories and MPXV samples are not readily available, we constructed a C9L genetic engineering model to simulate C9L mRNA expression in mammalian cells. The C9L gene of 9 MPXV strains was separately cloned into a vector plasmid and transiently transfected into HEK293FT cells (**Figure 4A**). To assess the effects of the C9L DNA G4 structure on its transcription, we evaluated the C9L mRNA level by RT-qPCR. As shown in **Figure S4**, the expression level of the C9L mRNA derived from 9 MPXV strains showed no obvious difference, indicating that the G4 structures of its genomic DNA cannot inhibit C9L mRNA transcription. It is because the C9L G4 motif is located in the non-templated strand of CDS region, instead of the promoter region that has stronger effects on transcription. Given that the C9L mRNA derived from 9 MPXV strains has similar expression level, it will be of interest to explore whether the ratio of G4 formation is related to the stability of the corresponding G4 motifs. Our group has developed the module assembly multifunctional probes assay (MAMPA) for the detection of RG4 in an individual gene in situ (*26*). Taking advantages of MAMPA, we detected the G4 structures of C9L mRNA derived from 9 MPXV strains. The MAMPA images of all the 9 variants of C9L mRNA showed typical G4 signals (**Figure 4B and Figure S5**). In contrast, when the G-tracts were G-A mutated, there was no observable G4 signal (**Figure S6**). These experiments provide evidence for the formation of RG4 structures in C9L mRNA in mammal cells. Moreover, as the stability of C9L RG4 in different MPXV strains increased, the average number of G4 fluorescent spots per cell also increased obviously (**Figure 4C**). Collectively, these data suggest that the in situ folding efficiency of the C9L RG4 variants was positively related to its stability.

**Figure 4.**
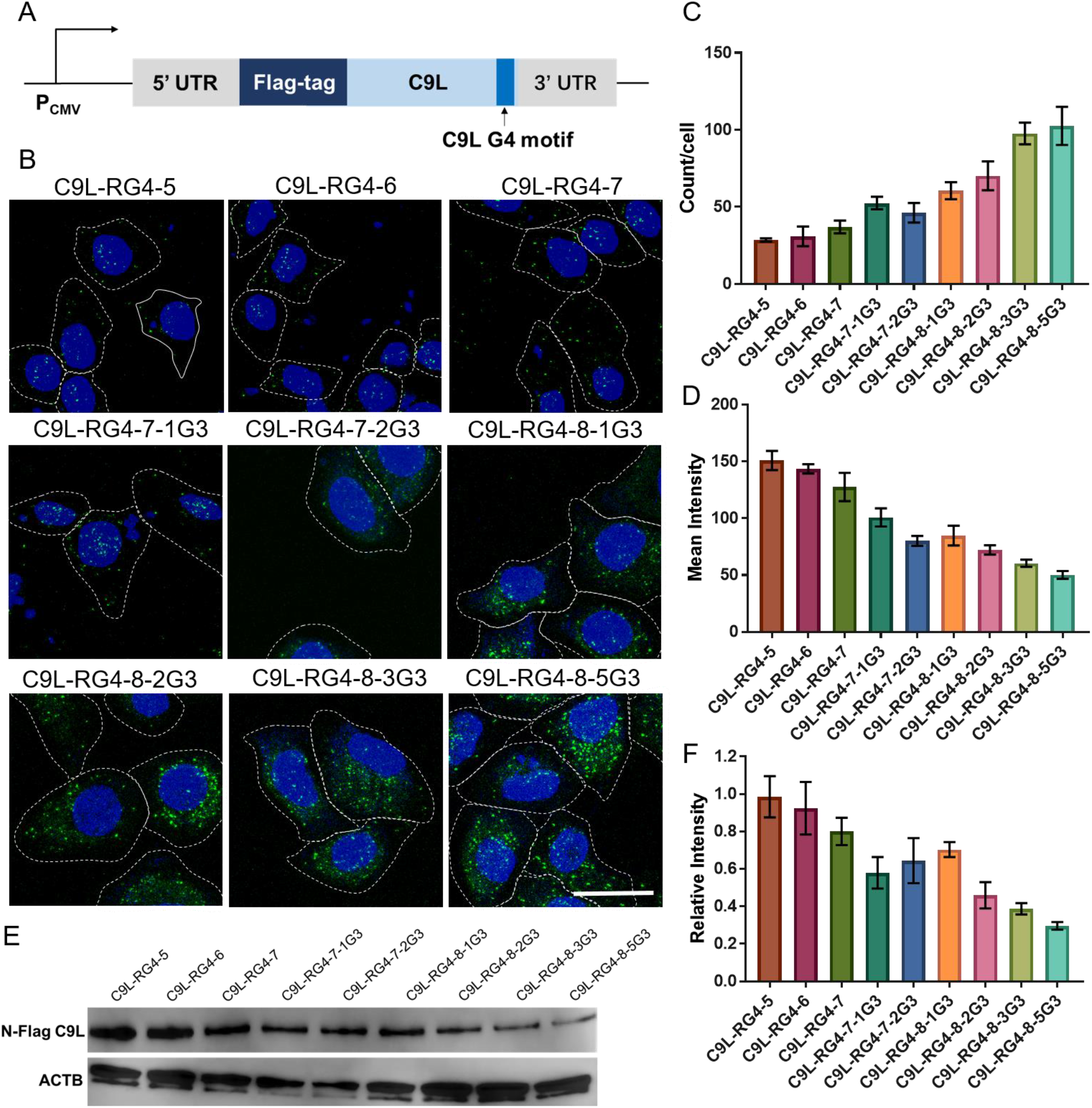
The evolution of C9L RG4 motifs reduces the formation of RG4 structures and increased C9L protein level in mammal cells. **A**) Schematic illustration of the C9L genetic engineering model to simulate C9L mRNA expression in mammalian cells. The whole-length CDS region of the C9L gene from 9 C9L variants was separately cloned into a mammal expression plasmid, and a Flag-tag was added to the N-terminal of CDS for immunoassay. **B**) MAMPA images of the 9 C9L RG4 motifs. The cell nuclei are shown in blue (DAPI), and the RCA particles appear as green spots (Alexa Fluor 488). Cells are outlined by white lines. Scale bar represents 25 μm. **C**) RCA particles per cell of each group in **B**. Data are from n=100 cells and presented as means ± SEM. **D**) Mean fluorescent intensity of N-flag C9L in each group of C9L RG4 variants. Fluorescence intensity was measured by flow cytometry. **E**) Western blotting analysis of N-flag C9L in each group of C9L RG4 variants. ACTB was used as a positive control. Relative amount of N-flag C9L in each group of C9L RG4 variants. N-flag C9L amount was normalized to the ACTB protein levels. Protein amount was measured by ImageJ software from the western blotting images. Data are shown as mean ± SEM of three independent experiments.

After validating the thermostability and in situ formation of the C9L RG4 variants, it will be of interest to further explore the relationships between the genomic evolution of MPXV and the generation of C9L RG4 variants with diverse stability. Strikingly, the C9L RG4 stability is generally lower in the West African clade than in the Central African clade (**Figure S1**). The strains of the West African clade after 2017 all contain the most unstable C9L-RG4-5 motif, while the strains of the Central African clade mainly contain the relative stable C9L-RG4-7-2G3. The RG4 of MPXV strains with similar evolutionary distances have similar stability. Additionally, the highly stable C9L RG4 motifs, C9L-RG4-8-2G3, C9L-RG4-8-3G3 and C9L-RG4-8-5G3, exist only in very few strains, indicating a low contagion. This data suggests that the evolution of MPXV tends to reduce the stability of C9L RG4 and favors the spread of the virus.

In general, the RG4 structure in the CDS region can inhibit the translation of the corresponding protein depending on the RG4 stability (*28*). Given that the C9L gene is related to the inhibition of innate immune response of the host, we infer that the evolution of C9L RG4 may regulate MPXV activity by increasing the C9L protein level. To assess the biological effects of the evolution of the C9L G4 motif, we focused on the translation efficiency of the corresponding protein. We evaluated the C9L protein expression in above mentioned C9L genetic engineering model. Using immunofluorescence and western blotting, we observed that the less stable C9L RG4 indeed increased the expression of C9L protein in living cells (**Figure 4D-4E**). These studies have led to the hypothesis that the increased transmission of MPXV is related to the evolution of C9L RG4. However, the understanding of the detail function of the C9L gene in the MPXV biology is very limited. Further studies that provide direct experimental support for the relationship between C9L RG4 and MPXV life cycle are necessary to advance our understanding of these findings.

## Conclusion

Since the eradication of smallpox in the world in 1980, monkeypox virus has become the most serious orthopox virus and keeps spreading across the world. The cause of the unusual MPXV epidemic remains elusive. We report that a RG4 motif located in the C9L gene CDS region and the C9L RG4 motif have evolved stepwise. The 9 C9L RG4 variants can fold into stable G4 structures in vitro. For the first time, we verified the formation of RG4 structures of the 9 C9L RG4 variants in living cells. Through phylogenetic analysis, we find that the evolution of MPXV generates increasingly unstable C9L RG4 motifs, and the current strains of the reported MKPV cases in 2022 all contain the most unstable C9L-RG4-5 motif. Next, we discover that the evolution of C9L RG4 motif resultes in an increased expression level of the corresponding protein. We inferr that this may cause the currently increased transmission of MPXV across the world. Taking together, these findings open a new avenue for the advance of MPXV biology, epidemiology, transmission, and pathogenesis. It is recommended that health authorities and researchers pay attention to the genomic evolution of MPXV.

## Acknowledgements

This work was financially supported by National Key Research and Development Program of China (No. 2021YFA1200104), National Natural Science Foundation of China (No. 22027807, No. 22034004, No. 21621003), the Strategic Priority Research Program of Chinese Academy of Sciences (No. XDB36000000), and Tsinghua University Spring Breeze Fund (No. 2020Z99CFZ019).

## Author Contributions

J.L. and X.T. conceived the idea and designed the experimental methodology. Y.D., X.T., D.H. and Q.Z. performed the experiments and collected the data. Y.D., X.T and J.L. analyzed the results and wrote the manuscript. J.L. supervised the entire project. Y.D. and X.T contributed equally to this work.

## Data Availability

All relevant data during the study are available from the corresponding authors upon request.

