## Supporting Information for "A peculiar evolutionary feature of monkeypox virus"

S2 **Supplementary Methods**

Synthesis of the oligos
Plasmid construction
Circular dichroism (CD) assay
Thioflavin T fluorescence assay
MAMPA imaging of the G-quadruplex in MKPV C9L mRNA
Cell culture and transfection
Flow cytometry
RT-qPCR
Western-blotting analysis

S6 **Supplementary Tables**

**Table S1** Oligonucleotide sequences used in this study
**Table S2** MKPV genomes used in this study.

S12 **Supplementary Figures**

**Figure S1** Detailed maximum likelihood phylogenetic tree of nucleotide sequences of the whole-genome of 177 MPXV strains.
**Figure S2** Thioflavin T fluorescent assay of the 9 C9L PQS oligos from different MKPV strains.
**Figure S3** CD spectra of the 9 C9L PQS oligos from different MKPV strains.
**Figure S4** RT-qPCR analysis of the C9L mRNA level of the 9 C9L variants.
**Figure S5** Zoom-in MAMPA images of the 9 C9L RG4 variants from different MKPV strains in living cells.
**Figure S6** MAMPA evaluation of the G4 structure formation of mutated C9L RG4 motifs in mammal cells.

### Supplementary Methods

#### Synthesis of the oligos

All the oligos used in this work (**Supplementary Table 1**) were purchased from Shanghai Sangon Biological Engineering Technology & Services Co., Ltd. (Shanghai, China), except the ID-Probe. ID-Probe was synthesized by amidation reaction between a 5'-$NH_2$, 3'-$N_3$ modified $A_{12}$ DNA strand and Carboxypyridostatin (CarboxyPDS, Sigma-Aldrich). 5'-$NH_2$, 3'-$N_3$ modified $A_{12}$ DNA strands was synthesized in a Mermade-12 Synthesiser (BioAutomation, USA) according to a previously reported protocol.

#### Plasmid construction

All the plasmids were prepared by inserting de novo synthesized C9L gene sequences from the 9 types of MPXV strains into the EcoRV site of pCDNA3.1(+) backbone plasmid. The sequence encoded 3X Flag-tag, GACTA CAAAG ACCAT GACGG TGATT ATAAA GATCA TGACA TCGAC TACAA GGATG ACGAT GACAA G, was added at the N-terminal of each C9L gene.

#### Circular dichroism (CD) assay

CD spectra and CD melting assays were conducted on a Chirascan plus spectropolarimeter with a temperature controller. Three scans were accumulated and automatically averaged. For the detection of CD melting curves, the signals were collected at a heating rate of 1 °C/min. The concentration of RNA was fixed at 2 μM.

#### Thioflavin T fluorescence assay

The fluorescence spectra were measured by an EnVision Multimodal Plate Readers (PerkinElmer) using an excitation wavelength of 442 nm. The concentration of ThT was fixed at 0.6 μM. The concentration of RNA was fixed at 0.6 μM.

#### MAMPA imaging of the G-quadruplex in MKPV C9L mRNA

Cells were cultured in 384-well glass bottom plates (Cellvis). Cells were fixed in 4% (w/v) paraformaldehyde (Beyotime) for 15 min at room temperature, washed twice with 1×DEPC-treated PBS (DEPC-PBS). Then, the cells were permeabilized for 5 min with 0.5% v/v Trion-X100 (Beijing Dingguo Changsheng Biotechnology) in 1×PBS. Next, 20 μL Probe-binding mixture [2×saline-sodium citrate buffer (SSC) (Ambion), 1 μM Amp-Probe, 1 μM ID-Probe,

1 mg/mL yeast tRNA (Solarbio), 5 mM DTT (Solarbio), 1 U/μL RiboLock RNase inhibitor (Thermo Scientific)] was added at 37 °C for 2 h. The sample was then washed three times using DEPC-PBS. Next, 20 μL padlock-binding mixture [2×SSC, 1 μM 5'-phosphorylated padlock DNA, 1 mg/mL yeast tRNA, 5 mM DTT, 1 U/μL RiboLock RNase inhibitor] overnight at 37 °C. Next, 10 μL padlock-cyclization mixture [1 ×ligase reaction buffer, 1 U/μL T4 DNA ligase (Thermo Scientific), 1 U/μL RiboLock RNase inhibitor] was added at 37 °C for 2 h. RCA reaction was then carried out in 10 μL RCA mixture [1×phi29 DNA polymerase reaction buffer, 0.4 U/μL phi29 DNA polymerase (Thermo Scientific), 3 mM dNTPs (Sangon), 1 U/μL of RiboLock RNase inhibitor] at 37 °C for 2 h. The hybridization of fluorescent probes was conducted in a 20 μL fluorescent probe-binding mixture [0.1 μM of fluorescent probe (Invitrogen), 2×SSC, 0.1 v/v formamide (Sangon), 10 ng/μL salmon sperm DNA (Solarbio)] at 37 °C for 30 min. After each step, the fixed cells were washed with 1X PBS (DEPC treated) for three times. After mounting with Fluoromount-G [containing 4′,6-diamidino-2-phenylindole (DAPI), SouthernBiotech], the samples were ready for imaging.

The samples were imaged by a Leica TCS SP5 inverted confocal microscope (Leica, Germany). All images were acquired with a 63×oil-immersion objective. The image size was 164 μm×164 μm, and the single pixel size was 160 nm×160 nm. The Alexa 488 dye was excited with an Ar laser (488 nm) and was detected with a 500-535 nm bandpass filter. The DAPI dye was excited with a HeNe543 laser (405 nm) and was detected with a 430-550 nm bandpass filter. Cy5 dye was excited with a HeNe633 laser and was detected with a 650-750 nm bandpass filter. Images were collected by layering with the step size of 0.2 μm to ensure that all RCA particles in cells were collected. The software Las AF version 2.6.3.8173 was used to stack with the method of maximum intensity project (MIP).

The RCA particles and the nuclei of cells in the fluorescent images were distinguished from the background by setting the threshold value using the software ImageJ version 1.46r. Then, the RCA particles and the nuclei of cells were merged by the software Image J, setting the RCA particles as green or red channel and the nuclei of cells as blue channel. The outline of cells was manually labeled from the bright field images and the numbers of RCA particles in a single cell were counted manually (n=100 cells in each group).

**Cell culture and transfection**

HEK293FT cells were obtained from National Infrastructure of Cell Line Resource (NICLR, China). The cell lines were checked free of mycoplasma contamination by PCR. Their species origins were confirmed with PCR. The identity of the cell line was authenticated with STR profiling (FBI, CODIS). All the results can be viewed on the website (http://cellresource.cn). HEK293FT cells were maintained in Dulbecco's modified eagle medium (DMEM, Gibco) supplemented with 10% FBS (Gibco) and 1% penicillin/streptomycin (Macklin) at 37 °C/5% $CO_2$.

HEK293FT cells cultured in 6-well plates were transfected at 80% confluence with 1 μg plasmid, 5 μL P3000™ Enhancer Reagent (Invitrogen) and 7 μL Lipofectamine® 3000 Reagent (Invitrogen). Further evaluation of the transfected cells were conducted after 48 h.

**Flow cytometry**

C9L gene transfected HEK293FT cells were cultured in a 6-well plate 2 days before test. After digested with 0.05% trypsin for 3 min, the cultured cells were centrifugated at 1,000 rpm for 3 minutes and the supernatant was removed. Cells were fixed in 4% paraformaldehyde and permeabilized in 0.5% Triton X-100. After blocking for 30 min at 37 °C in 5% BSA/1XPBS, cells were incubated with the DYKDDDDK Tag Antibody (1:200 dilution, Cell Signaling Technology, #2368). After three rinses in 1XPBS, cells were incubated with Mouse Anti-rabbit IgG/FITC Antibody (1:1,000 dilution, #bs-0295M-FITC, Bioss) for 1 h at 37 °C. Then, cells were resuspended by 300 μL complete medium and the solution was placed in flow cytometry tube for detection. The fluorescence intensity of cells was measured by BD LSRFortessa™ cell analyzer. The forward scattered light (FSC) and lateral scattered light (SSC) parameters were used to determine the range of single cells, and then the single cell population was analyzed. The fluorescence intensity of cells in the FITC channel were detected to measure the expression level of C9L in cells.

**RT-qPCR**

Total RNAs of cells were extracted by RNAprep pure Cell / Bacteria Kit (Tiangen) according to the manufacturer's protocol. The concentration and the purify of RNAs were measured by a Nanodrop 2000 UV-vis spectrophotometer. Total RNAs were reverse transcribed with random primers (Yeasen) and RevertAid Reverse Transcriptase (Thermo Scientific) under the manufacturer's protocol. qPCR was conducted using Hieff UNICON qPCR SYBR Green Master Mix (Yeasen) under the manufacturer's protocol. The primers used in qPCR were listed in **Table S1**. Ct values were measured by a Bio-Rad CFX96 (Bio-Rad) instrument and were averaged from three replicate measurements. Expression levels of the studied genes were calculated by $2^{-\Delta\Delta Ct}$ method and the house-keeping *ACTB* was used as internal controls.

**Western-blotting analysis**

Proteins in cells were extracted by Tissue or Cell Total Protein Extration Kit (Sangon) according to the manufacturer's protocol. Protein lysates were separated on SDS-PAGEs and blotted on a Polyvinylidene Fluoride (PVDF) membrane (Beyotime). After blocking free binding sites with 5% milk powder in 1× TBS-T, membrane was incubated with the first antibody for 1 h at room temperature under the constant agitation. After three times 7 min washing with Washing Buffer (Beyotime), membrane was incubated with HRP-conjugated second antibody for 1 h at room temperature followed by another three washing steps. Signals were detected by chemiluminescence on a Gel Doc XR+ Gel Documentation System (Biorad).

For the detection of N-Flag C9L protein, the first antibody is DYKDDDDK Tag Antibody (1:1000 dilution, Cell Signaling Technology, #2368), and the second antibody is Goat Anti-rabbit IgG/HRP Antibody (1:5000, Bioss, #bs-0295G-HRP). For the detection of ACTB protein, the first antibody is Beta Actin Polyclonal Antibody (1:5000, Yeasen, #30102ES40), and the second antibody is Goat Anti-rabbit IgG/HRP Antibody (1:5000, Bioss, #bs-0295G-HRP).

### Supplementary Tables

**Table S1** Oligonucleotide sequences used in this study

| Name | Sequences (5'-3') | Description |
|---|---|---|
| C9L-RG4-5 | GGAAUAUAUGGGAUGGAAUGGAAUGGUAAAUAAUUUGAAA | C9L RG4 oligos of different MKPV strains |
| C9L-RG4-6 | GGAAUAUAUGGGAUGGAAUGGAAUGGAAUGGUAAAUAAUUU | |
| C9L-RG4-7 | GGAAUAUAUGGAAUGGAAUGGAAUGGAAUGGAAUGGUAAAU | |
| C9L-RG4-7-1G3 | GGAAUAUAUGGGAUGGAAUGGAAUGGAAUGGAAUGGUAAAU | |
| C9L-RG4-7-2G3 | GGAAUAUAUGGAAUGGGAUGGGAUGGAAUGGAAUGGUAAAU | |
| C9L-RG4-8-1G3 | GGAAUAUAUGGGAUGGAAUGGAAUGGAAUGGAAUGGAAUGG | |
| C9L-RG4-8-2G3 | GGAAUAUAUGGGAUGGAAUGGGAUGGAAUGGAAUGGAAUGG | |
| C9L-RG4-8-3G3 | GGAAUAUAUGGAAUGGGAUGGGAUGGGAUGGAAUGGAAUGG | |
| C9L-RG4-8-5G3 | GGAAUAUAUGGAAUGGGAUGGGAUGGGAUGGGAUGGGAUG G | |
| MUT-C9L-RG4-5 | AAAAUAUAUAAAAUAAAAUAAAAUAAUAAAUAAUUUAAAA | G-A mutated C9L RG4 oligos of different MKPV strains |
| MUT-C9L-RG4-6 | AAAAUAUAUAAAAUAAAAUAAAAUAAAAUAAUAAAUAAUUU | |
| MUT-C9L-RG4-7 | AAAAUAUAUAAAAUAAAAUAAAAUAAAAUAAAAUAAUAAAU | |
| MUT-C9L-RG4-7-1G3 | AAAAUAUAUAAAAUAAAAUAAAAUAAAAUAAAAUAAUAAAU | |
| MUT-C9L-RG4-7-2G3 | AAAAUAUAUAAAAUAAAAUAAAAUAAAAUAAAAUAAUAAAU | |
| MUT-C9L-RG4-8-1G3 | AAAAUAUAUAAAAUAAAAUAAAAUAAAAUAAAAUAAAAUAA | |
| MUT-C9L-RG4-8-2G3 | AAAAUAUAUAAAAUAAAAUAAAAUAAAAUAAAAUAAAAUAA | |
| MUT-C9L-RG4-8-3G3 | AAAAUAUAUAAAAUAAAAUAAAAUAAAAUAAAAUAAAAUAA | |
| MUT-C9L-RG4-8-5G3 | AAAAUAUAUAAAAUAAAAUAAAAUAAAAUAAAAUAAAAUAA | |
| C9L-F | TCCGATGAATAGCCCCAGAC | qPCR primers |
| C9L-R | GGAACCAACGCTCAACAGATG | |
| ID-Probe | /CarboxyPDS/AAAAAAAAAAAA/$N_3$-C/ | DNA probes used in MAMPA |
| Amp-Probe | /DBCO/CCATGAATAAGTGCGATTAT GCTAGCTAGC | |
| Padlock Probe | /Phos/TTTTTTCCC AACTATACAACATACTACCTCA CTT GCTAGCTAGC ATAATCGCACTTATTCATGG TTTTTTTT | |

| Fluorescent Probe | /FAM/AACTATACAACATACTACCTCA |
|---|---|

**Table S2** MKPV genomes used in this study.

Accession, GenBank Accession number. Country, the country in which the MKPV strain was discovered. Year, the time at which the MKPV strain was discovered. G-Tracts, the number of G-tracts. The “XG3” postfix means the number of the G-tract with 3 G-bases. G4 sequence, the sequence of the G4 motif.

| Accession | Country | Year | G-Tracts | G4 Sequence |
|---|---|---|---|---|
| MG693723.1 | Nigeria | 2017 | 5 | GGAATATATGGGATGGAATGGAATGGTAAATAATTTTGAAA |
| MG693724.1 | Nigeria | 2017 | 5 | GGAATATATGGGATGGAATGGAATGGTAAATAATTTTGAAA |
| MG693725.1 | Nigeria | 2017 | 5 | GGAATATATGGGATGGAATGGAATGGTAAATAATTTTGAAA |
| MK783027.1 | Nigeria | 2017 | 5 | GGAATATATGGGATGGAATGGAATGGTAAATAATTTTGAAA |
| MK783028.1 | Nigeria | 2017 | 5 | GGAATATATGGGATGGAATGGAATGGTAAATAATTTTGAAA |
| MK783029.1 | Nigeria | 2017 | 5 | GGAATATATGGGATGGAATGGAATGGTAAATAATTTTGAAA |
| MK783030.1 | Nigeria | 2017 | 5 | GGAATATATGGGATGGAATGGAATGGTAAATAATTTTGAAA |
| MK783031.1 | Nigeria | 2017 | 5 | GGAATATATGGGATGGAATGGAATGGTAAATAATTTTGAAA |
| MK783032.1 | Nigeria | 2017 | 5 | GGAATATATGGGATGGAATGGAATGGTAAATAATTTTGAAA |
| MK783033.1 | Nigeria | 2017 | 5 | GGAATATATGGGATGGAATGGAATGGTAAATAATTTTGAAA |
| MN648051.1 | Israel | 2019 | 5 | GGAATATATGGGATGGAATGGAATGGTAAATAATTTTGAAA |
| MT903337.1 | Nigeria | 2020 | 5 | GGAATATATGGGATGGAATGGAATGGTAAATAATTTTGAAA |
| MT903338.1 | Nigeria | 2020 | 5 | GGAATATATGGGATGGAATGGAATGGTAAATAATTTTGAAA |
| MT903339.1 | Nigeria | 2020 | 5 | GGAATATATGGGATGGAATGGAATGGTAAATAATTTTGAAA |
| MT903340.1 | Nigeria | 2020 | 5 | GGAATATATGGGATGGAATGGAATGGTAAATAATTTTGAAA |
| MT903341.1 | Nigeria | 2020 | 5 | GGAATATATGGGATGGAATGGAATGGTAAATAATTTTGAAA |
| MT903342.1 | Singapore | 2020 | 5 | GGAATATATGGGATGGAATGGAATGGTAAATAATTTTGAAA |
| MT903343.1 | UK | 2020 | 5 | GGAATATATGGGATGGAATGGAATGGTAAATAATTTTGAAA |
| MT903344.1 | UK | 2020 | 5 | GGAATATATGGGATGGAATGGAATGGTAAATAATTTTGAAA |
| MT903345.1 | UK | 2020 | 5 | GGAATATATGGGATGGAATGGAATGGTAAATAATTTTGAAA |
| ON563414.2 | USA | 2022 | 5 | GGAATATATGGGATGGAATGGAATGGTAAATAATTTTGAAA |
| ON568298.1 | Germany | 2022 | 5 | GGAATATATGGGATGGAATGGAATGGTAAATAATTTTGAAA |
| ON585029.1 | Portugal | 2022 | 5 | GGAATATATGGGATGGAATGGAATGGTAAATAATTTTGAAA |
| ON585030.1 | Portugal | 2022 | 5 | GGAATATATGGGATGGAATGGAATGGTAAATAATTTTGAAA |
| ON585031.1 | Portugal | 2022 | 5 | GGAATATATGGGATGGAATGGAATGGTAAATAATTTTGAAA |
| ON585032.1 | Portugal | 2022 | 5 | GGAATATATGGGATGGAATGGAATGGTAAATAATTTTGAAA |
| ON585033.1 | Portugal | 2022 | 5 | GGAATATATGGGATGGAATGGAATGGTAAATAATTTTGAAA |
| ON585034.1 | Portugal | 2022 | 5 | GGAATATATGGGATGGAATGGAATGGTAAATAATTTTGAAA |
| ON585035.1 | Portugal | 2022 | 5 | GGAATATATGGGATGGAATGGAATGGTAAATAATTTTGAAA |
| ON585036.1 | Portugal | 2022 | 5 | GGAATATATGGGATGGAATGGAATGGTAAATAATTTTGAAA |
| ON585037.1 | Portugal | 2022 | 5 | GGAATATATGGGATGGAATGGAATGGTAAATAATTTTGAAA |

| | | | | |
|---|---|---|---|---|
| ON585038.1 | Portugal | 2022 | 5 | GGAATATATGGGATGGAATGGAATGGTAAATAATTTTGAAA |
| ON595760.1 | Switzerland | 2022 | 5 | GGAATATATGGGATGGAATGGAATGGTAAATAATTTTGAAA |
| ON602722.1 | France | 2022 | 5 | GGAATATATGGGATGGAATGGAATGGTAAATAATTTTGAAA |
| ON619835.2 | UK | 2022 | 5 | GGAATATATGGGATGGAATGGAATGGTAAATAATTTTGAAA |
| ON619836.2 | UK | 2022 | 5 | GGAATATATGGGATGGAATGGAATGGTAAATAATTTTGAAA |
| ON619837.2 | UK | 2022 | 5 | GGAATATATGGGATGGAATGGAATGGTAAATAATTTTGAAA |
| ON619838.2 | UK | 2022 | 5 | GGAATATATGGGATGGAATGGAATGGTAAATAATTTTGAAA |
| ON736420.1 | Canada | 2022 | 5 | GGAATATATGGGATGGAATGGAATGGTAAATAATTTTGAAA |
| ON602722.2 | France | 2022 | 5 | GGAATATATGGGATGGAATGGAATGGTAAATAATTTTGAAA |
| ON720848.1 | Spain | 2022 | 5 | GGAATATATGGGATGGAATGGAATGGTAAATAATTTTGAAA |
| ON720849.1 | Spain | 2022 | 5 | GGAATATATGGGATGGAATGGAATGGTAAATAATTTTGAAA |
| ON694329.1 | Germany | 2022 | 5 | GGAATATATGGGATGGAATGGAATGGTAAATAATTTTGAAA |
| ON694330.1 | Germany | 2022 | 5 | GGAATATATGGGATGGAATGGAATGGTAAATAATTTTGAAA |
| ON694331.1 | Germany | 2022 | 5 | GGAATATATGGGATGGAATGGAATGGTAAATAATTTTGAAA |
| ON694332.1 | Germany | 2022 | 5 | GGAATATATGGGATGGAATGGAATGGTAAATAATTTTGAAA |
| ON694333.1 | Germany | 2022 | 5 | GGAATATATGGGATGGAATGGAATGGTAAATAATTTTGAAA |
| ON694334.1 | Germany | 2022 | 5 | GGAATATATGGGATGGAATGGAATGGTAAATAATTTTGAAA |
| ON694335.1 | Germany | 2022 | 5 | GGAATATATGGGATGGAATGGAATGGTAAATAATTTTGAAA |
| ON694336.1 | Germany | 2022 | 5 | GGAATATATGGGATGGAATGGAATGGTAAATAATTTTGAAA |
| ON694337.1 | Germany | 2022 | 5 | GGAATATATGGGATGGAATGGAATGGTAAATAATTTTGAAA |
| ON694338.1 | Germany | 2022 | 5 | GGAATATATGGGATGGAATGGAATGGTAAATAATTTTGAAA |
| ON694339.1 | Germany | 2022 | 5 | GGAATATATGGGATGGAATGGAATGGTAAATAATTTTGAAA |
| ON694340.1 | Germany | 2022 | 5 | GGAATATATGGGATGGAATGGAATGGTAAATAATTTTGAAA |
| ON694341.1 | Germany | 2022 | 5 | GGAATATATGGGATGGAATGGAATGGTAAATAATTTTGAAA |
| ON694342.1 | Germany | 2022 | 5 | GGAATATATGGGATGGAATGGAATGGTAAATAATTTTGAAA |
| ON682263.2 | Germany | 2022 | 5 | GGAATATATGGGATGGAATGGAATGGTAAATAATTTTGAAA |
| ON682264.2 | Germany | 2022 | 5 | GGAATATATGGGATGGAATGGAATGGTAAATAATTTTGAAA |
| ON682265.2 | Germany | 2022 | 5 | GGAATATATGGGATGGAATGGAATGGTAAATAATTTTGAAA |
| ON682266.1 | Germany | 2022 | 5 | GGAATATATGGGATGGAATGGAATGGTAAATAATTTTGAAA |
| ON682267.1 | Germany | 2022 | 5 | GGAATATATGGGATGGAATGGAATGGTAAATAATTTTGAAA |
| ON682268.1 | Germany | 2022 | 5 | GGAATATATGGGATGGAATGGAATGGTAAATAATTTTGAAA |
| ON682269.2 | Germany | 2022 | 5 | GGAATATATGGGATGGAATGGAATGGTAAATAATTTTGAAA |
| ON682270.1 | Germany | 2022 | 5 | GGAATATATGGGATGGAATGGAATGGTAAATAATTTTGAAA |
| ON674051.1 | USA | 2022 | 5 | GGAATATATGGGATGGAATGGAATGGTAAATAATTTTGAAA |
| ON675438.1 | USA | 2022 | 5 | GGAATATATGGGATGGAATGGAATGGTAAATAATTTTGAAA |
| ON676703.1 | USA | 2022 | 5 | GGAATATATGGGATGGAATGGAATGGTAAATAATTTTGAAA |
| ON676704.1 | USA | 2022 | 5 | GGAATATATGGGATGGAATGGAATGGTAAATAATTTTGAAA |
| ON676705.1 | USA | 2022 | 5 | GGAATATATGGGATGGAATGGAATGGTAAATAATTTTGAAA |
| ON676706.1 | USA | 2022 | 5 | GGAATATATGGGATGGAATGGAATGGTAAATAATTTTGAAA |
| ON676707.1 | USA | 2022 | 5 | GGAATATATGGGATGGAATGGAATGGTAAATAATTTTGAAA |
| ON676708.1 | USA | 2022 | 5 | GGAATATATGGGATGGAATGGAATGGTAAATAATTTTGAAA |
| ON609725.2 | Slovenia | 2022 | 5 | GGAATATATGGGATGGAATGGAATGGTAAATAATTTTGAAA |
| ON622722.2 | France | 2022 | 5 | GGAATATATGGGATGGAATGGAATGGTAAATAATTTTGAAA |

| | | | | |
|---|---|---|---|---|
| ON637938.1 | Germany | 2022 | 5 | GGAATATATGGGATGGAATGGAATGGTAAATAATTTTGAAA |
| ON637939.1 | Germany | 2022 | 5 | GGAATATATGGGATGGAATGGAATGGTAAATAATTTTGAAA |
| ON644344.1 | Italy | 2022 | 5 | GGAATATATGGGATGGAATGGAATGGTAAATAATTTTGAAA |
| ON645312.1 | UK | 2022 | 5 | GGAATATATGGGATGGAATGGAATGGTAAATAATTTTGAAA |
| ON649708.1 | Portugal | 2022 | 5 | GGAATATATGGGATGGAATGGAATGGTAAATAATTTTGAAA |
| ON649709.1 | Portugal | 2022 | 5 | GGAATATATGGGATGGAATGGAATGGTAAATAATTTTGAAA |
| ON649710.1 | Portugal | 2022 | 5 | GGAATATATGGGATGGAATGGAATGGTAAATAATTTTGAAA |
| ON649711.1 | Portugal | 2022 | 5 | GGAATATATGGGATGGAATGGAATGGTAAATAATTTTGAAA |
| ON649712.1 | Portugal | 2022 | 5 | GGAATATATGGGATGGAATGGAATGGTAAATAATTTTGAAA |
| ON649713.1 | Portugal | 2022 | 5 | GGAATATATGGGATGGAATGGAATGGTAAATAATTTTGAAA |
| ON649714.1 | Portugal | 2022 | 5 | GGAATATATGGGATGGAATGGAATGGTAAATAATTTTGAAA |
| ON649715.1 | Portugal | 2022 | 5 | GGAATATATGGGATGGAATGGAATGGTAAATAATTTTGAAA |
| ON649716.1 | Portugal | 2022 | 5 | GGAATATATGGGATGGAATGGAATGGTAAATAATTTTGAAA |
| ON649717.1 | Portugal | 2022 | 5 | GGAATATATGGGATGGAATGGAATGGTAAATAATTTTGAAA |
| ON649718.1 | Portugal | 2022 | 5 | GGAATATATGGGATGGAATGGAATGGTAAATAATTTTGAAA |
| ON649719.1 | Portugal | 2022 | 5 | GGAATATATGGGATGGAATGGAATGGTAAATAATTTTGAAA |
| ON649720.1 | Portugal | 2022 | 5 | GGAATATATGGGATGGAATGGAATGGTAAATAATTTTGAAA |
| ON649721.1 | Portugal | 2022 | 5 | GGAATATATGGGATGGAATGGAATGGTAAATAATTTTGAAA |
| ON649722.1 | Portugal | 2022 | 5 | GGAATATATGGGATGGAATGGAATGGTAAATAATTTTGAAA |
| ON649723.1 | Portugal | 2022 | 5 | GGAATATATGGGATGGAATGGAATGGTAAATAATTTTGAAA |
| ON649724.1 | Portugal | 2022 | 5 | GGAATATATGGGATGGAATGGAATGGTAAATAATTTTGAAA |
| ON649725.1 | Portugal | 2022 | 5 | GGAATATATGGGATGGAATGGAATGGTAAATAATTTTGAAA |
| ON595760.2 | Switzerland | 2022 | 5 | GGAATATATGGGATGGAATGGAATGGTAAATAATTTTGAAA |
| ON631963.1 | Australia | 2022 | 5 | GGAATATATGGGATGGAATGGAATGGTAAATAATTTTGAAA |
| ON563414.3 | USA | 2022 | 5 | GGAATATATGGGATGGAATGGAATGGTAAATAATTTTGAAA |
| ON622720.1 | Switzerland | 2022 | 5 | GGAATATATGGGATGGAATGGAATGGTAAATAATTTTGAAA |
| ON631241.1 | Slovenia | 2022 | 5 | GGAATATATGGGATGGAATGGAATGGTAAATAATTTTGAAA |
| ON627808.1 | USA | 2022 | 5 | GGAATATATGGGATGGAATGGAATGGTAAATAATTTTGAAA |
| ON622712.1 | Belgium | 2022 | 5 | GGAATATATGGGATGGAATGGAATGGTAAATAATTTTGAAA |
| ON622713.1 | Belgium | 2022 | 5 | GGAATATATGGGATGGAATGGAATGGTAAATAATTTTGAAA |
| ON622718.1 | Spain | 2022 | 5 | GGAATATATGGGATGGAATGGAATGGTAAATAATTTTGAAA |
| ON622721.1 | Italy | 2022 | 5 | GGAATATATGGGATGGAATGGAATGGTAAATAATTTTGAAA |
| ON614676.1 | Italy | 2022 | 5 | GGAATATATGGGATGGAATGGAATGGTAAATAATTTTGAAA |
| ON615424.1 | Netherlands | 2022 | 5 | GGAATATATGGGATGGAATGGAATGGTAAATAATTTTGAAA |
| KJ642617.1 | Nigeria | 1971 | 6 | GGAATATATGGGATGGAATGGAATGGAATGGTAAATAATTT |
| DQ011155.1 | Zaire | 1978 | 6 | GGAATATATGGAATGGGATGGAATGGAATGGTAAATAATTT |
| HM172544.1 | Zaire | 1979 | 6 | GGAATATATGGAATGGGATGGAATGGAATGGTAAATAATTT |
| KC257459.1 | Sudan | 2005 | 6 | GGAATATATGGAATGGGATGGAATGGAATGGTAAATAATTT |
| KC257460.1 | Sudan | 2005 | 6 | GGAATATATGGAATGGGATGGAATGGAATGGTAAATAATTT |
| ON649879.1 | Israel | 2022 | 6 | GGAATATATGGGATGGAATGGAATGGTAAATAATTTTGAAA |
| KJ642614.1 | Netherlands | 1965 | 7 | GGAATATATGGAATGGAATGGAATGGAATGGAATGGTAAAT |
| AY741551.1 | Sierra Leone | 1970 | 7-1G3 | GGAATATATGGGATGGAATGGAATGGAATGGAATGGTAAAT |

| | | | | |
|---|---|---|---|---|
| DQ011156.1 | Liberia | 1970 | 7-1G3 | GGAATATATGGGATGGAATGGAATGGAATGGAATGGTAAAT |
| KJ642615.1 | Nigeria | 1978 | 7-1G3 | GGAATATATGGGATGGAATGGAATGGAATGGAATGGTAAAT |
| KJ136820.1 | Cote d'Ivoire | 2012 | 7-1G3 | GGAATATATGGGATGGAATGGAATGGAATGGAATGGTAAAT |
| KJ642613.1 | Zaire | 1970 | 7-2G3 | GGAATATATGGAATGGGATGGGATGGAATGGAATGGTAAAT |
| KJ642619.1 | Gabon | 1987 | 7-2G3 | GGAATATATGGAATGGGATGGGATGGAATGGAATGGTAAAT |
| MN702453.1 | Central African Republic | 2001 | 7-2G3 | GGAATATATGGAATGGGATGGGATGGAATGGAATGGTAAAT |
| DQ011154.1 | Congo | 2003 | 7-2G3 | GGAATATATGGAATGGGATGGGATGGAATGGAATGGTAAAT |
| JX878407.1 | Congo | 2006 | 7-2G3 | GGAATATATGGAATGGGATGGGATGGAATGGAATGGTAAAT |
| JX878409.1 | Congo | 2006 | 7-2G3 | GGAATATATGGAATGGGATGGGATGGAATGGAATGGTAAAT |
| JX878410.1 | Congo | 2006 | 7-2G3 | GGAATATATGGAATGGGATGGGATGGAATGGAATGGTAAAT |
| JX878411.1 | Congo | 2006 | 7-2G3 | GGAATATATGGAATGGGATGGGATGGAATGGAATGGTAAAT |
| JX878412.1 | Congo | 2006 | 7-2G3 | GGAATATATGGAATGGGATGGGATGGAATGGAATGGTAAAT |
| JX878413.1 | Congo | 2006 | 7-2G3 | GGAATATATGGAATGGGATGGGATGGAATGGAATGGTAAAT |
| JX878414.1 | Congo | 2006 | 7-2G3 | GGAATATATGGAATGGGATGGGATGGAATGGAATGGTAAAT |
| JX878415.1 | Congo | 2006 | 7-2G3 | GGAATATATGGAATGGGATGGGATGGAATGGAATGGTAAAT |
| JX878416.1 | Congo | 2006 | 7-2G3 | GGAATATATGGAATGGGATGGGATGGAATGGAATGGTAAAT |
| JX878421.1 | Congo | 2007 | 7-2G3 | GGAATATATGGAATGGGATGGGATGGAATGGAATGGTAAAT |
| JX878422.1 | Congo | 2007 | 7-2G3 | GGAATATATGGAATGGGATGGGATGGAATGGAATGGTAAAT |
| JX878423.1 | Congo | 2007 | 7-2G3 | GGAATATATGGAATGGGATGGGATGGAATGGAATGGTAAAT |
| JX878424.1 | Congo | 2007 | 7-2G3 | GGAATATATGGAATGGGATGGGATGGAATGGAATGGTAAAT |
| JX878425.1 | Congo | 2007 | 7-2G3 | GGAATATATGGAATGGGATGGGATGGAATGGAATGGTAAAT |
| JX878427.1 | Congo | 2007 | 7-2G3 | GGAATATATGGAATGGGATGGGATGGAATGGAATGGTAAAT |
| JX878428.1 | Congo | 2007 | 7-2G3 | GGAATATATGGAATGGGATGGGATGGAATGGAATGGTAAAT |
| JX878429.1 | Congo | 2007 | 7-2G3 | GGAATATATGGAATGGGATGGGATGGAATGGAATGGTAAAT |
| KP849469.1 | Congo | 2008 | 7-2G3 | GGAATATATGGAATGGGATGGGATGGAATGGAATGGTAAAT |
| MN702452.1 | Central African Republic | 2010 | 7-2G3 | GGAATATATGGAATGGGATGGGATGGAATGGAATGGTAAAT |
| MT724770.1 | Congo | 2014 | 7-2G3 | GGAATATATGGAATGGGATGGGATGGAATGGAATGGTAAAT |
| MT724771.1 | Congo | 2014 | 7-2G3 | GGAATATATGGAATGGGATGGGATGGAATGGAATGGTAAAT |
| MN702448.1 | Central African Republic | 2016 | 7-2G3 | GGAATATATGGAATGGGATGGGATGGAATGGAATGGTAAAT |
| MN702449.1 | Central African Republic | 2016 | 7-2G3 | GGAATATATGGAATGGGATGGGATGGAATGGAATGGTAAAT |
| MN702450.1 | Central African Republic | 2016 | 7-2G3 | GGAATATATGGAATGGGATGGGATGGAATGGAATGGTAAAT |

| | | | | |
|---|---|---|---|---|
| MN702444.1 | Central African Republic | 2017 | 7-2G3 | GGAATATATGGAATGGGATGGGATGGAATGGAATGGTAAAT |
| MN702445.1 | Central African Republic | 2017 | 7-2G3 | GGAATATATGGAATGGGATGGGATGGAATGGAATGGTAAAT |
| MN702451.1 | Central African Republic | 2017 | 7-2G3 | GGAATATATGGAATGGGATGGGATGGAATGGAATGGTAAAT |
| MN702446.1 | Central African Republic | 2018 | 7-2G3 | GGAATATATGGAATGGGATGGGATGGAATGGAATGGTAAAT |
| MN702447.1 | Central African Republic | 2018 | 7-2G3 | GGAATATATGGAATGGGATGGGATGGAATGGAATGGTAAAT |
| AY753185.1 | Denmark | 1958 | 8-1G3 | GGAATATATGGGATGGAATGGAATGGAATGGAATGGAATGG |
| AY603973.1 | USA | 1971 | 8-1G3 | GGAATATATGGGATGGAATGGAATGGAATGGAATGGAATGG |
| KP849470.1 | Cote d'Ivoire | 1971 | 8-1G3 | GGAATATATGGGATGGAATGGAATGGAATGGAATGGAATGG |
| MN346692.1 | Ivory Coast | 2017 | 8-1G3 | GGAATATATGGGATGGAATGGAATGGAATGGAATGGAATGG |
| MN346693.1 | Ivory Coast | 2017 | 8-1G3 | GGAATATATGGGATGGAATGGAATGGAATGGAATGGAATGG |
| MN346696.1 | Ivory Coast | 2017 | 8-1G3 | GGAATATATGGGATGGAATGGAATGGAATGGAATGGAATGG |
| MN346697.1 | Ivory Coast | 2017 | 8-1G3 | GGAATATATGGGATGGAATGGAATGGAATGGAATGGAATGG |
| MN346698.1 | Ivory Coast | 2017 | 8-1G3 | GGAATATATGGGATGGAATGGAATGGAATGGAATGGAATGG |
| MN346701.1 | Ivory Coast | 2017 | 8-1G3 | GGAATATATGGGATGGAATGGAATGGAATGGAATGGAATGG |
| MN346702.1 | Ivory Coast | 2018 | 8-1G3 | GGAATATATGGGATGGAATGGAATGGAATGGAATGGAATGG |
| KP849471.1 | Zaire | 1979 | 8-2G3 | GGAATATATGGAATGGGATGGAATGGGATGGAATGGAATGG |
| KJ642612.1 | Zaire | 1986 | 8-2G3 | GGAATATATGGAATGGGATGGGATGGAATGGAATGGAATGG |
| KJ642618.1 | Cameroon | 1989 | 8-2G3 | GGAATATATGGAATGGGATGGGATGGAATGGAATGGAATGG |
| DQ011153.1 | USA | 2003 | 8-2G3 | GGAATATATGGGATGGAATGGGATGGAATGGAATGGAATGG |
| DQ011157.1 | USA | 2003 | 8-2G3 | GGAATATATGGGATGGAATGGGATGGAATGGAATGGAATGG |
| MT903346.1 | USA | 2004 | 8-2G3 | GGAATATATGGGATGGAATGGGATGGAATGGAATGGAATGG |
| MT903347.1 | USA | 2005 | 8-2G3 | GGAATATATGGGATGGAATGGGATGGAATGGAATGGAATGG |
| MT903348.1 | USA | 2006 | 8-2G3 | GGAATATATGGGATGGAATGGGATGGAATGGAATGGAATGG |
| JX878408.1 | Congo | 2009 | 8-3G3 | GGAATATATGGAATGGGATGGGATGGGATGGAATGGAATGG |
| JX878417.1 | Congo | 2009 | 8-3G3 | GGAATATATGGAATGGGATGGGATGGGATGGAATGGAATGG |
| JX878418.1 | Congo | 2009 | 8-3G3 | GGAATATATGGAATGGGATGGGATGGGATGGAATGGAATGG |
| JX878419.1 | Congo | 2009 | 8-3G3 | GGAATATATGGAATGGGATGGGATGGGATGGAATGGAATGG |
| JX878420.1 | Congo | 2009 | 8-3G3 | GGAATATATGGAATGGGATGGGATGGGATGGAATGGAATGG |
| JX878426.1 | Congo | 2009 | 8-3G3 | GGAATATATGGAATGGGATGGGATGGGATGGAATGGAATGG |
| AF380138.1 | Zaire | 1996 | 8-5G3 | GGAATATATGGAATGGGATGGGATGGGATGGGATGGGATGG |

### Supplementary Figures

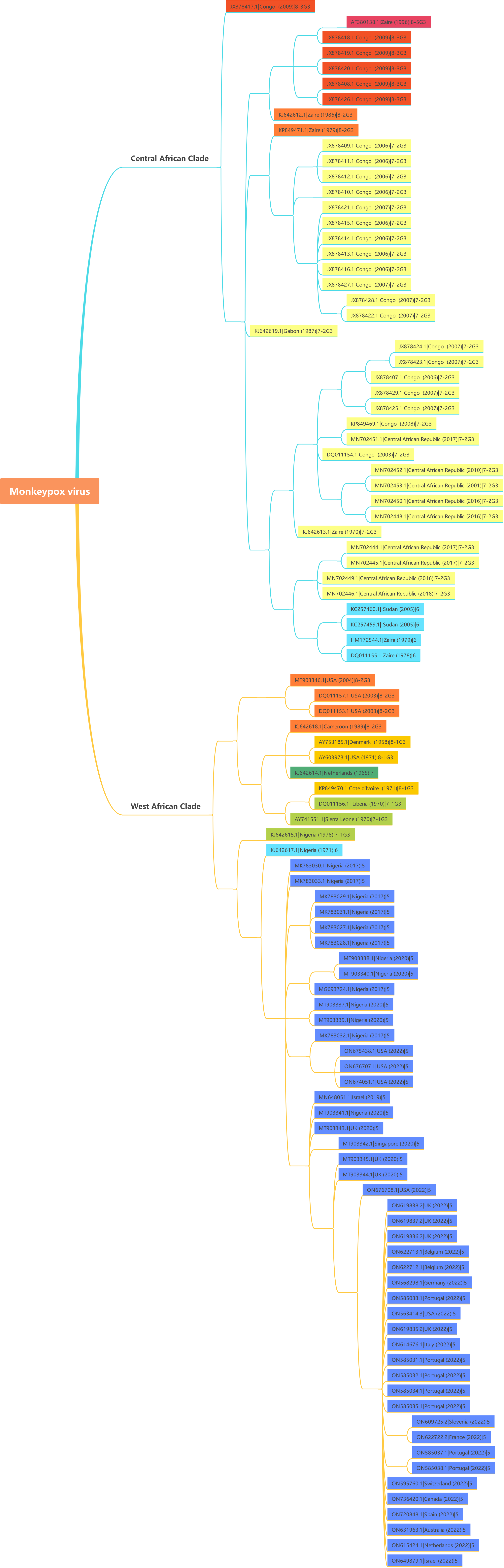

Monkeypox virus
Central African Clade
JX878417.1|Congo (2009)|8-3G3
AF380138.1|Zaire (1996)|8-5G3
JX878418.1|Congo (2009)|8-3G3
JX878419.1|Congo (2009)|8-3G3
JX878420.1|Congo (2009)|8-3G3
JX878408.1|Congo (2009)|8-3G3
JX878426.1|Congo (2009)|8-3G3
KJ642612.1|Zaire (1986)|8-2G3
KP849471.1|Zaire (1979)|8-2G3
JX878409.1|Congo (2006)|7-2G3
JX878411.1|Congo (2006)|7-2G3
JX878412.1|Congo (2006)|7-2G3
JX878410.1|Congo (2006)|7-2G3
JX878421.1|Congo (2007)|7-2G3
JX878415.1|Congo (2006)|7-2G3
JX878414.1|Congo (2006)|7-2G3
JX878413.1|Congo (2006)|7-2G3
JX878416.1|Congo (2006)|7-2G3
JX878427.1|Congo (2007)|7-2G3
JX878428.1|Congo (2007)|7-2G3
JX878422.1|Congo (2007)|7-2G3
KJ642619.1|Gabon (1987)|7-2G3
JX878424.1|Congo (2007)|7-2G3
JX878423.1|Congo (2007)|7-2G3
JX878407.1|Congo (2006)|7-2G3
JX878429.1|Congo (2007)|7-2G3
JX878425.1|Congo (2007)|7-2G3
KP849469.1|Congo (2008)|7-2G3
MN702451.1|Central African Republic (2017)|7-2G3
DQ011154.1|Congo (2003)|7-2G3
MN702452.1|Central African Republic (2010)|7-2G3
MN702453.1|Central African Republic (2001)|7-2G3
MN702450.1|Central African Republic (2016)|7-2G3
MN702448.1|Central African Republic (2016)|7-2G3
KJ642613.1|Zaire (1970)|7-2G3
MN702444.1|Central African Republic (2017)|7-2G3
MN702445.1|Central African Republic (2017)|7-2G3
MN702449.1|Central African Republic (2016)|7-2G3
MN702446.1|Central African Republic (2018)|7-2G3
KC257460.1| Sudan (2005)|6
KC257459.1| Sudan (2005)|6
HM172544.1|Zaire (1979)|6
DQ011155.1|Zaire (1978)|6
West African Clade
MT903346.1|USA (2004)|8-2G3
DQ011157.1|USA (2003)|8-2G3
DQ011153.1|USA (2003)|8-2G3
KJ642618.1|Cameroon (1989)|8-2G3
AY753185.1|Denmark (1958)|8-1G3
AY603973.1|USA (1971)|8-1G3
KJ642614.1|Netherlands (1965)|7
KP849470.1|Cote d'Ivoire (1971)|8-1G3
DQ011156.1| Liberia (1970)|7-1G3
AY741551.1|Sierra Leone (1970)|7-1G3
KJ642615.1|Nigeria (1978)|7-1G3
KJ642617.1|Nigeria (1971)|6
MK783030.1|Nigeria (2017)|5
MK783033.1|Nigeria (2017)|5
MK783029.1|Nigeria (2017)|5
MK783031.1|Nigeria (2017)|5
MK783027.1|Nigeria (2017)|5
MK783028.1|Nigeria (2017)|5
MT903338.1|Nigeria (2020)|5
MT903340.1|Nigeria (2020)|5
MG693724.1|Nigeria (2017)|5
MT903337.1|Nigeria (2020)|5
MT903339.1|Nigeria (2020)|5
MK783032.1|Nigeria (2017)|5
ON675438.1|USA (2022)|5
ON676707.1|USA (2022)|5
ON674051.1|USA (2022)|5
MN648051.1|Israel (2019)|5
MT903341.1|Nigeria (2020)|5
MT903343.1|UK (2020)|5
MT903342.1|Singapore (2020)|5
MT903345.1|UK (2020)|5
MT903344.1|UK (2020)|5
ON676708.1|USA (2022)|5
ON619838.2|UK (2022)|5
ON619837.2|UK (2022)|5
ON619836.2|UK (2022)|5
ON622713.1|Belgium (2022)|5
ON622712.1|Belgium (2022)|5
ON568298.1|Germany (2022)|5
ON585033.1|Portugal (2022)|5
ON563414.3|USA (2022)|5
ON619835.2|UK (2022)|5
ON614676.1|Italy (2022)|5
ON585031.1|Portugal (2022)|5
ON585032.1|Portugal (2022)|5
ON585034.1|Portugal (2022)|5
ON585035.1|Portugal (2022)|5
ON609725.2|Slovenia (2022)|5
ON622722.2|France (2022)|5
ON585037.1|Portugal (2022)|5
ON585038.1|Portugal (2022)|5
ON595760.1|Switzerland (2022)|5
ON736420.1|Canada (2022)|5
ON720848.1|Spain (2022)|5
ON631963.1|Australia (2022)|5
ON615424.1|Netherlands (2022)|5
ON649879.1|Israel (2022)|5

**Figure S1** Detailed maximum likelihood phylogenetic tree of nucleotide sequences of the whole-genome of 177 MPXV strains. The type of C9L RG4 motif of each strain was noted in the end of the strain names and labelled with different colors.

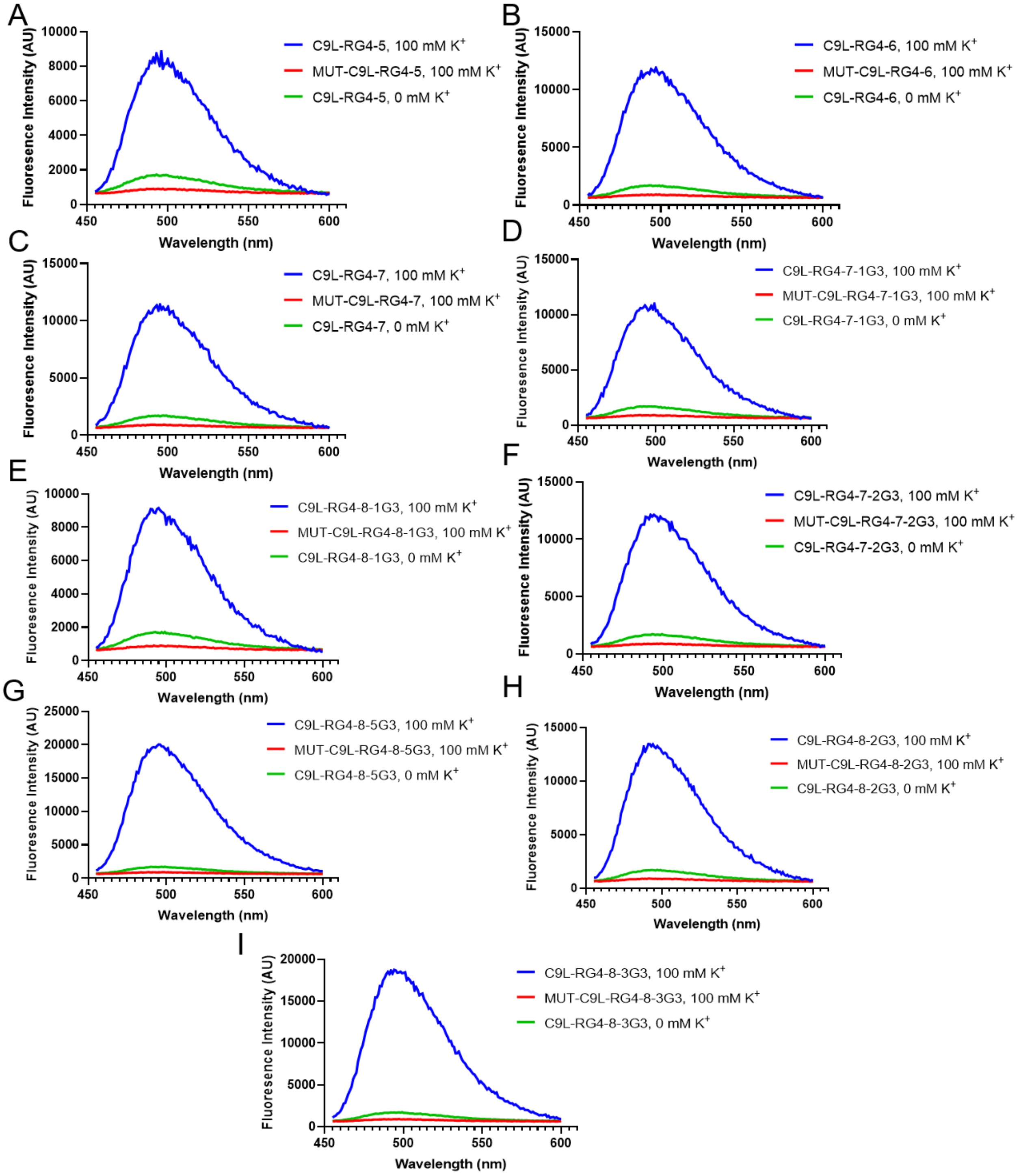

**Figure S2** Thioflavin T fluorescent assay of the 9 C9L PQS oligos from different MKPV strains. The mutated C9L PQS oligos and the assays in 0 mM $K^+$ were negative controls in each group.

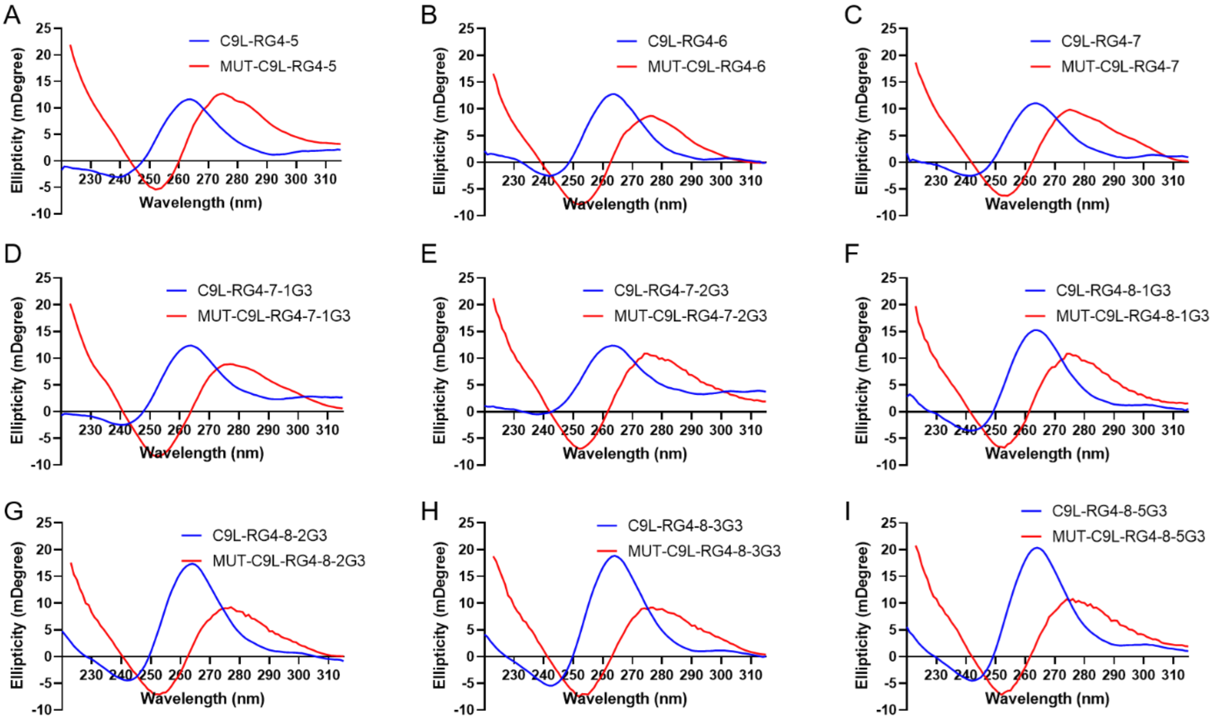

**Figure S3** CD spectra of the 9 C9L PQS oligos from different MKPV strains. The mutated C9L PQS oligos were negative controls in each group.

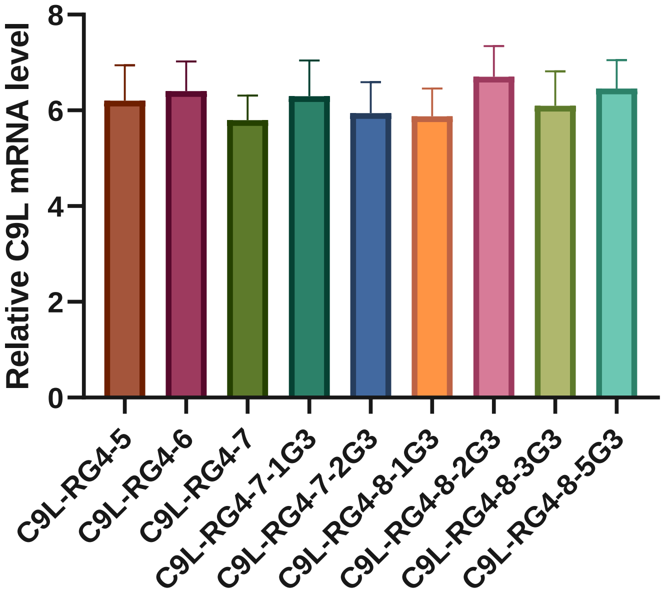

**Figure S4** RT-qPCR analysis of the C9L mRNA level of the 9 C9L variants. Data are shown as mean ± SEM of three independent experiments.

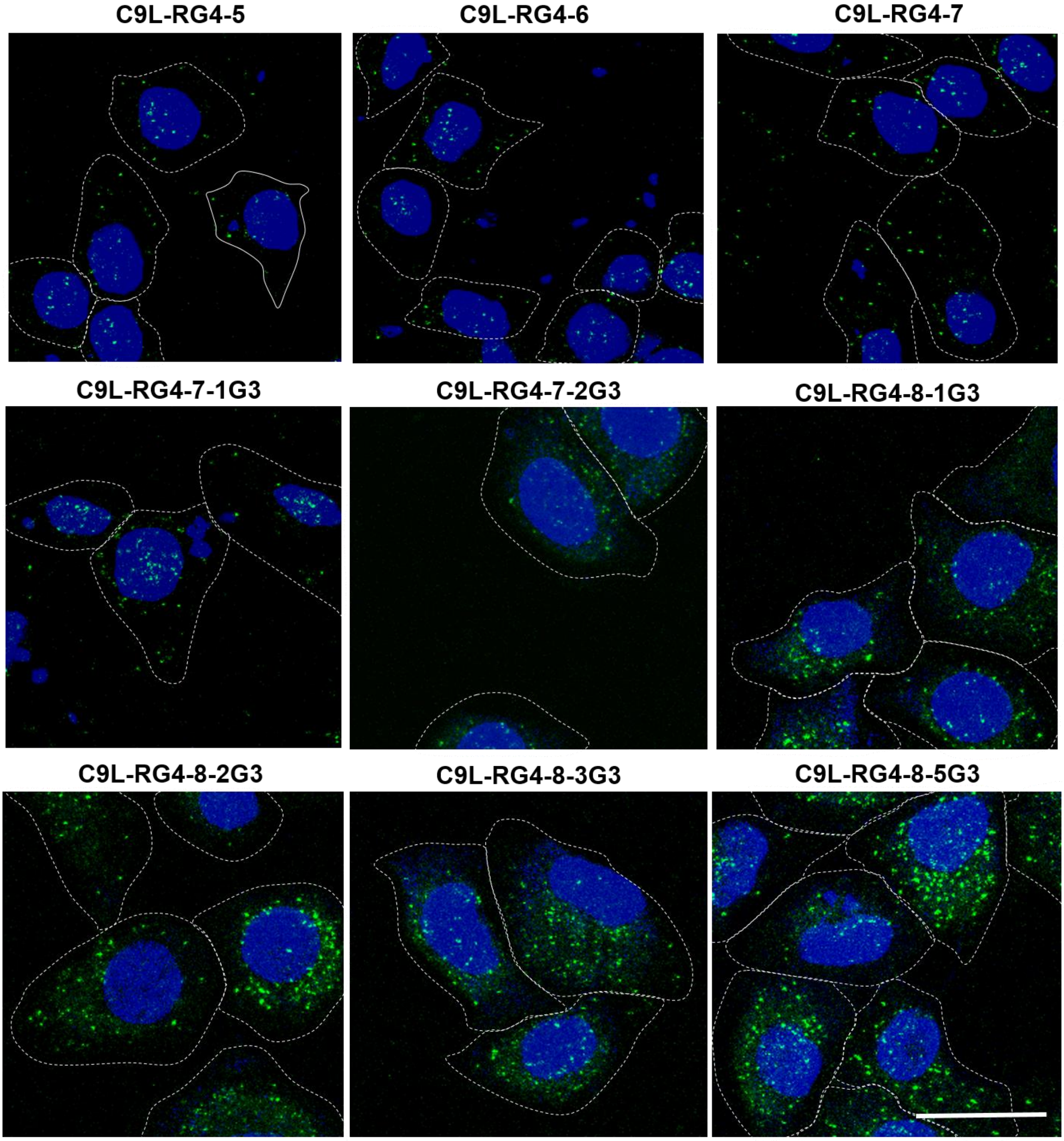

**Figure S5** Zoom-in MAMPA images of the 9 C9L RG4 variants from different MKPV strains in living cells.

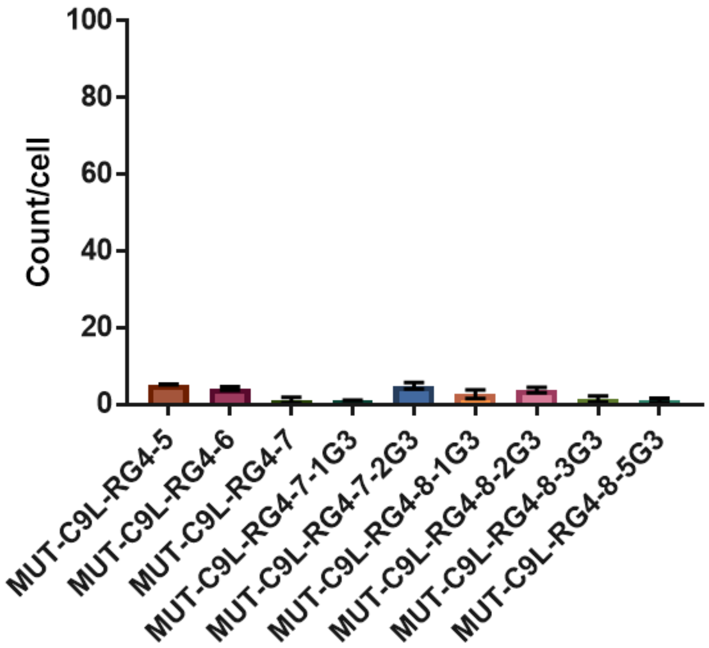

**Figure S6** MAMPA evaluation of the G4 structure formation of mutated C9L RG4 motifs in mammal cells. RCA particles per cell of each group were counted from MAMPA images. Data are from n=100 cells and presented as means ± SEM.
